# *In silico* simulations reveal molecular mechanism of uranyl ion toxicity towards DNA-binding domain of PARP-1 protein

**DOI:** 10.1101/2023.06.02.543417

**Authors:** Egor S. Bulavko, Dmitry N. Ivankov

## Abstract

The molecular toxicity of uranyl ion (UO_2_^2+^) in living cells is mainly conditioned by its high affinity to both native and potential metal-binding sites frequently occurring in biomolecules structure. Recent advances in computational and experimental research shed light on the structural properties and functional impacts of uranyl binding to proteins, organic ligands, nucleic acids and their complexes. In the present work, we report the results of the theoretical investigation of the uranyl-mediated loss of DNA-binding activity of PARP-1, eukaryotic enzyme that participates in DNA reparation, cell differentiation, induction of inflammation, etc. Latest experimental studies showed that uranyl ion directly interacts with its DNA-binding subdomains – zinc fingers Zn1 and Zn2, – and changes their tertiary structure. Here, we propose an atomistic mechanism underlying this process and compute the free energy change along the suggested pathway to prove its relevance. According to the results of our QM/MM simulations of Zn2-UO_2_^2+^ complex, uranyl ion replaces zinc in its native binding site, but the corresponding state is destroyed because of the following spontaneous internal hydrolysis of the U–Cys162 coordination bond. Although the enthalpy of hydrolysis is +2.8 kcal/mol, the final value of the free energy of the reaction constitutes -0.6 kcal/mol, due to structure loosening evidenced by solvation and configuration thermodynamic properties calculated using GIST- and MIST-based trajectory processing techniques. The subsequent reorganization of the binding site includes association of uranyl ion with the Glu190/Asp191 acidic cluster and significant perturbations in the domain’s tertiary structure, which further decreases the free energy of the non-functional state by 6.8 kcal/mol. The disruption of the DNA-binding interface revealed in our computational simulations is consistent with previous experimental findings and appears to be associated with the loss of the Zn2 affinity for nucleic acids.

## Introduction

Uranium (U) is a poisonous heavy metal that is primarily of concern due to inherent radioactivity. However, under natural conditions, its chemical toxicity can pose even a greater risk [1]. Uranyl ion (UO_2_^2+^), the most stable soluble form of uranium, showed an ability to interfere with numerous biochemical processes [2]. For instance, uranyl substitutes iron in the natural metal-binding site of hemoglobin, considerably reducing the amount of its oxy-form in blood [3]. Uranyl also disrupts the formation of the *Cyt c / Cyt b*_*5*_ complex, which participates in the process of cell respiration [4]. By unraveling the intricate molecular interactions and pathways involved in these and other uranyl-mediated toxic effects, we can gain insights into their detrimental impacts on biological systems. This knowledge can provide a solid foundation for studying the toxicological aspects of uranyl ion exposure, identifying potential biomarkers of toxicity and developing targeted therapeutic strategies.

Although numerous experimental and computational studies have focused on the ability of uranyl ion to induce conformational changes in individual proteins and disrupt their complexes [5, 6], there has been relatively limited research on the structural mechanisms related to the uranyl-mediated disruption of DNA-protein interactions. It has long been known that uranium poisoning is associated with DNA repair deficiency [7, 8]. In 2016, Cooper with coauthors showed that uranyl ion is a direct inhibitor of Poly [ADP-ribose] polymerase 1 (PARP-1), eukaryotic enzyme which, among other activities, recognizes single- and double-stranded DNA breaks and initiates various repair mechanisms [9]. The DNA-binding domain (DBD) of PARP-1 consists of three zinc finger motifs, Zn1, Zn2, and Zn3 [10]. The first two are highly homologous and make a major contribution to the recognition of DNA damages (Fig. 1). The authors hypothesized that Zn1 and Zn2 domains are the main targets of uranium, as far as PARP-1 exposure to uranyl acetate caused zinc loss from the protein [9]. Zhou and coauthors confirmed this in 2021 [11]. Working with an isolated peptide corresponding to the Zn1 zinc-binding site, they found that uranyl ion destroys its native tertiary structure, which was maintained both in the absence of any metal and upon incubation with zinc salts.

**Figure 1.**
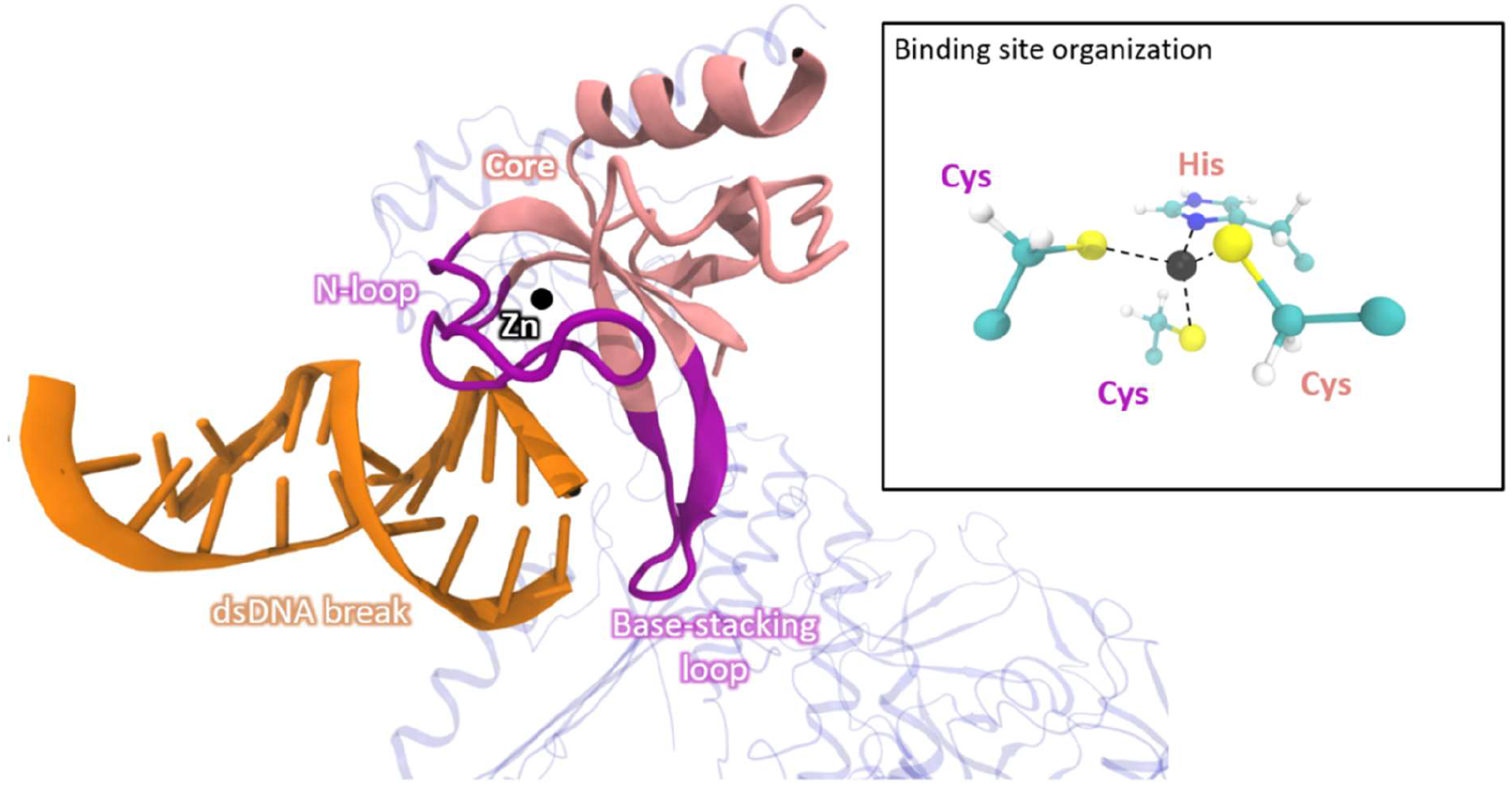
Detailed structure of Zn1 zinc finger bound to double-stranded DNA break in the PARP-1 environment (PDB ID 4DQY). N-terminal loop (N-loop) recognizes major groove of DNA fragment, and base-stacking loop interacts with nucleotides’ aromatic rings. Zinc ion binding site is composed of two cysteines located within N-loop and one histidine and one cysteine from protein’s core.

Here, we applied a wide arsenal of computational biochemistry techniques to investigate the structural properties of uranyl ion interaction with PARP-1 zinc fingers and find a possible molecular pathway resulting in the decrease of the domains’ affinity to nucleic acids. The chemical transformations involving uranyl ion in protein’s metal-binding sites and corresponding free energy changes were studied using molecular dynamics with *ab initio* type QM/MM potentials, which was proven to be accepted practice in this field [12-14]. To explore the conformational space of the zinc finger/uranyl complex, we performed classical molecular dynamics simulations using one of the standard protein force fields, which was additionally parametrized for accounting for uranyl ion coordinated by particular amino acids side chains. Finally, we compared the thermodynamic stabilities of the discovered conformational states by computing their relative solvation and configurational enthalpies and entropies.

## Models and Methods

An atomistic molecular model for computational experiments was built based on the X-ray crystalline structure of the Zn2 zinc finger of human PARP-1 (PDB ID 3ODC) [15]. We dissolved the protein in a rectangular 7.1 × 6.8 × 6.3 nm^3^ water box and added sodium and chloride ions to neutralize the system and make NaCl concentration equal to 150 mM. CHARMM36 force field [16-18] and TIP3P water model [19] were utilized to generate topology that was then converted from CHARMM to GROMACS format. All actions described above were performed in VMD program (version 1.9.4a53) [20], including psfgen and topogromacs [21] plugins for building and manipulating topology. Metal-binding sites search in the starting model was performed using MIB2 web server (http://bioinfo.cmu.edu.tw/MIB2) [22].

Classical molecular dynamics simulations were conducted in GROMACS 2021.6 (mixed precision) program package [23]. Typical simulation setup implied utilization of V-rescale thermostat [24] (T = 300 K, time constant 1 ps) with separated temperature coupling for solute and solvent molecules, isotropic Parrinello-Rahman barostat (P = 1 atm, time constant 5 ps, compressibility 1 atm^-1^) and h-bonds constraints with LINCS [25] and SHAKE [26] algorithms for solute and water, respectively. Coulomb and VdW cutoffs were set to 1.2 nm, long-range electrostatics was computed with Particle Mesh Ewald (PME) method [27], and the overall integration time step was set to 2 fs.

To study the electronic structure and bonding properties of complexes with uranyl and zinc ions we performed pure quantum chemistry calculations. More than 10 four- and five-coordinated complexes varying in coordination sphere composition were explored. The set of possible addends included hydroxyl, acetate (CH_3_COO^−^) and methanethiolate (CH_3_S^−^) ions, as well as water molecules. We conducted geometry optimizations at the DFT/PBE0/D3BJ/ZORA level of theory [28-30]. Pople-style basis set 6-311++G** [31, 32] was utilized for light atoms and zinc, while SARC-ZORA-TZVP [33] – for uranium. Additionally, we applied the implicit CPCM solvation model [34]. We performed these calculations in ORCA (version 5.0.1) program package [35]. Final wave functions were postprocessed using Multiwfn (version 3.8) program [36]. We calculated bond lengths, Mayer bond orders, Mulliken charges and Intrinsic bond strength indexes (IBSI).

The chemical transformations occurring in the protein’s metal-binding sites and corresponding free energy changes were studied throughout QM/MM molecular dynamics in combination with bias potentials (Umbrella Sampling, US) [37]. The quantum subsystem was composed of uranyl ion, surrounding amino acids’ side chains (namely, Cys125, Cys128, His159, Cys162 at the first stage and Cys125, Cys128, Glu190 at the second) along with one-two catalytic water molecules. Forces acting in quantum subsystem were described at DFT/PBE0/D3BJ level of theory with 6-31G** basis for light atoms and LANL2DZ [38] for uranium. Metal’s non-valence electrons were replaced by HayWadt effective core potential (ECP) [39] for further reduction of computational load. QM-MM interactions were dealt with in terms of electrostatic embedding; Mulliken atomic charges were computed for the quantum subsystem at every step of molecular dynamics. The boundary effects were resolved using the link atom scheme. The classical part was described with CHARMM36 force field, solvent – with TIP3P water model. We performed QM/MM calculations with the help of NAMD (version 2.14) – ORCA program interface [40]. Hybrid molecular dynamics setup implied usage of Langevin thermostat (T = 300 K, damping coefficient 5 ps^-1^), Nosé-Hoover Langevin piston barostat (P = 1 atm, oscillation period 200 fs, damping scale 100 fs), integration step 1 fs and the absence of any bond constraints. The non-bonded cutoff was set to 1.2 nm, and long-range electrostatics was computed using PME. Weighted Histograms Analysis (WHAM) [41] and Umbrella Integration (UI) [42] methods were used to process the results of Umbrella Sampling calculations.

We additionally evaluated the enthalpy of chemical reactions by comparing bare electronic energies of the QM subsystem in initial and final states. From 5 ps simulations of reactant and product we randomly selected twenty snapshots and performed geometry optimizations of quantum subsystem in the presence of point-charge correction conditioned by 1730 carefully selected solute and solvent atoms and absolute coordinate restraints applied to link hydrogens. The difference between average energies of optimal geometries was then compared with those resulted from the Umbrella Sampling simulation.

To parametrize uranium coordination complexes that correspond to the observed metastable states in terms of classical force field, we utilized a standard protocol implemented as the Force Field toolkit (ffTK) plugin [43] in VMD. Restrained electrostatic potential (RESP) partial charges were assigned to atoms, and VdW parameters for uranyl were obtained from one of the previous studies [44].

We used the GIST (Grid inhomogeneous solvation theory) methodology [45, 46] implemented in AMBER CPPTRAJ program [47] to estimate the solvation enthalpy and entropy of different protein states. A 5000 ns trajectory was computed for each state and clustered using gromacs cluster module (cluster method – gromos, cutoff – 0.5 nm). The most representative structure from each cluster underwent additional 500 ns simulations (resulted in 50,000 snapshots in total) with protein atoms positions restrained. These trajectories were analyzed using GIST. ParmEd python library [48] was utilized to convert topology files to format suitable for CPPTRAJ. The GIST grid size and center were chosen to include the whole protein and a 10 Å solvent buffer zone. Grid spacing was set to 0.5 Å for water entropy and 1.0 Å for water energy calculations. Multiple GIST runs were performed, gradually increasing the frames analyzed until the target thermodynamic properties values reached a plateau. Water-water energy was calculated from a pairwise interaction matrix [46]. The total solvation free energy included translational and orientational water entropy terms, water-solute and water-water energy. For protein states represented by multiple structures, solvation energy was calculated by weighted averaging.

Configurational enthalpy in the particular state was estimated as a sum of internal electrostatic and VdW interactions. We omitted all bonded terms because they are commonly relaxed, on average. A hundred random snapshots was taken from each cluster and proceeded by the g_mmpbsa program [49] to evaluate the target properties. The PARENT program [50] was utilized to calculate configurational entropy by MIST (Mutual information spanning tree) method. 1500 ns trajectories (with coordinates written every ps) were analyzed. The number of bins was set to 50 for 1D histograms and to 2500 – for 2D ones.

## Results and Discussion

### Structural properties of uranyl-protein complex

To begin with, we performed a structure-based search of metal-binding sites on the Zn2 zinc finger (PDB ID 3ODC) surface using MIB2 web tool [22]. Aside from the native zinc-binding site, we identified two new potential sites (Fig. 2A). Both consist of neighboring acidic residues, specifically Asp and Glu, which form a cluster capable of accommodating up to four coordination covalent bonds (CCBs). One potential site occupies the terminus of the base-stacking loop, while the other is located within one of the core’s alpha helices. Since exposure of full-length PARP-1 and its DNA-binding domain to uranyl salts resulted in zinc loss [9, 11], we chose the native binding site for our investigation.

**Figure 2.**
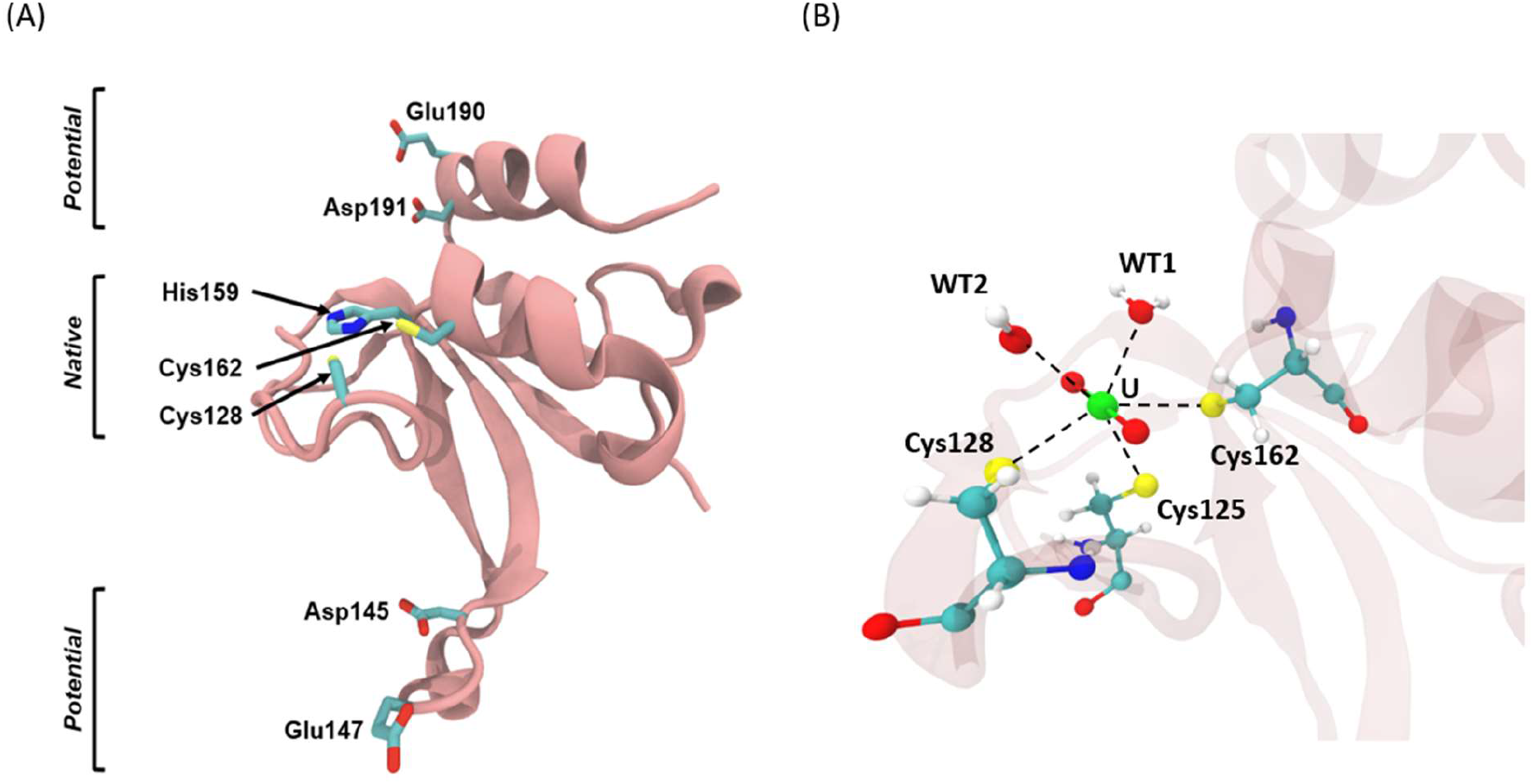
(A) Zinc finger metal-binding sites identified by MIB2 server [22]. Cys125 residue, which is a part of the native site, is not shown. (B) Uranyl ion binding pose in native site. WT stands for water.

Naturally, four amino acids side chains coordinate zinc in its binding site. Namely, the N-loop contains two cysteine residues, which, along with histidine and cysteine from the protein’s core, form a tetrahedral environment for the zinc ion (Fig. 1). We performed quantum chemistry calculations (for details see “Models and Methods”) and found that the zinc-sulfur bond in the complex has a characteristic length of 2.3 Å, and the zinc-nitrogen bond – 1.8 Å. The average bond orders for Zn-S and Zn-N bonds equaled 0.84 and 0.91, respectively. We also found that the zinc charge decreased from +2.0 to +0.9 upon binding to protein. This suggests that the covalent contribution in the binding process is predominant, indicating a strong and stable complex formation. As a result, the disordered loop is tightly bound to the protein’s core, which is essential for achieving high affinity to DNA [15].

It is well known that uranyl ion prefers a flat coordination sphere consisting of four, five or six ligands. Furthermore, due to its strong acceptor properties, it has a higher affinity for charged ligands compared to neutral ones. Consequently, we anticipated significant perturbations upon the substitution of zinc with uranium in the native binding site. QM/MM molecular dynamics simulation revealed that in this case His159 leaves the coordination environment, and the remaining cysteine residues along with two water molecules form an irregular pentagon, lying in a plane perpendicular to the O-U-O axis (Fig. 2B). The bond between uranium and sulfur atoms in the complex were found to be longer and weaker compared to the corresponding zinc-sulfur bond, with an average length of 2.75 Å, and the bond order equal to 0.65. The distances between uranium and oxygen of the coordinated water molecules are comparable to the uranium-sulfur distances, equaling 2.71 Å. However, the corresponding bond order are relatively small, being 0.28. Only a tiny charge, -0.13, is transferred from water to uranium indicating a lower covalent contribution to the U-H_2_O bond.

### Coordination environment determines the complex stability

The precise positioning of the N-loop is crucial for DNA recognition, and zinc plays a critical role in maintaining its proper orientation [15]. Therefore, we aimed to investigate the extent to which uranyl can assume the connection between N-loop and protein’s core.

We analyzed how the composition of coordination environment affects the overall stability of the complex. We investigated over 10 complexes and evaluated four bond characteristics: bond length, Mayer order, IBSI (intrinsic bond strength index), and transferred Mullliken charge. Mayer bond order was shown to be a valuable metric of complexation effectiveness [51], so we were primarily interested in its correlation with the coordination sphere content.

The results of our calculations are given in Table S1. Figure 3 reveals two important findings. Firstly, as hydroxyl anions are added into the coordination sphere, there is a gradual decrease in the average U-S bond order. This suggests that the presence of small charged ligands weakens the bond strength between uranium and sulfur atoms. Secondly, the U-S bond order in five-coordinated complexes, as a general trend, is lower with respect to that in four-coordinated complexes, which highlights the significant role played by steric factors in determining the stability of such complexes. Thus, if the coordination sphere consists of two CH_3_S^−^ ions and two water molecules, the U-S bond order is 0.85, and it becomes as low as 0.36 if the environment contains three CH_3_S- and two OH-groups. Interestingly, neither U-H_2_O nor U-OH interaction properties are changed upon variation of the complex composition. Here, the deviations of bond lengths and orders from complex to complex do not exceed 11% (Table S1). Thus, our findings demonstrate that under specific conditions, the uranium-sulfur bonds can weaken and undergo dissociation.

**Figure 3.**
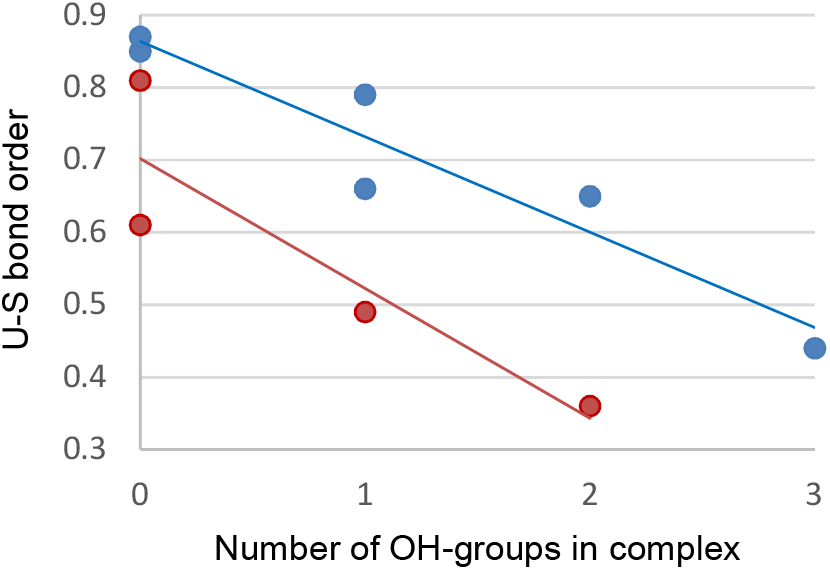
Correlation between Mayer bond order of uranium-sulfur bond and ligand content, namely the number of OH- and CH_3_S-groups, in uranyl ion complexes (blue stands for two Cys-like addends, red – for three).

### Cysteine 162 discoordination leads to the loss of zinc finger tertiary structure

The U-Cys162 bond was of particular importance in our study, as its disruption could result in the loss of connectivity between the zinc finger domains (see Fig. 2B) and potentially lead to a change in orientation of the N-loop, which is undesirable for DNA recognition [15]. As demonstrated earlier, the initial coordination environment composition includes three cysteines, two water molecules, and no hydroxyl ions. This configuration corresponds to a relatively high bond order of 0.78. Replacing one water molecule with hydroxyl anion would yield a complex with the formula [UO_2_(SCH_3_)_3_(OH)(H_2_O)]^2-^. Despite quantum chemistry calculations indicated its instability, we estimated the U-S bond order for this modified complex to be lower than 0.49. The process of water deprotonation could occur concurrently with cysteine dissociation within a process called internal hydrolysis:

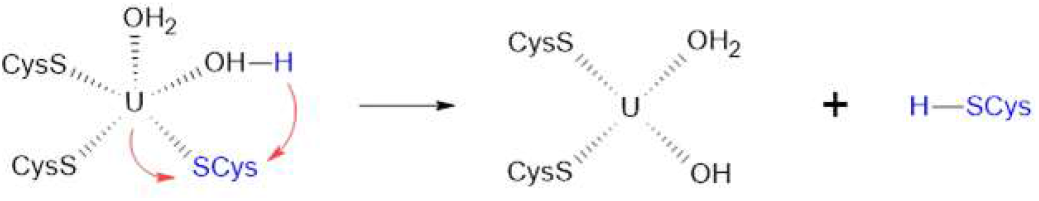

It implies the transfer of a proton from water to cysteine, which further facilitates the breaking of the uranium-sulfur bond, since the uranyl ion exhibits stronger acidic properties than hydrogen sulfide and its derivatives [52].

To evaluate the energy of chemical reactions, several methods are widely utilized. One of the most common is Umbrella Sampling (US), which accounts for not only enthalpy but entropy of a process as well. However, it is unable to separate these two energy terms, so that, to estimate the hydrolysis enthalpy alone, we used the protocol described in “Models and Methods” section.

The free energy cross-section profile of the hydrolysis reaction obtained from Umbrella Sampling calculations is presented in supplementary materials (Fig. S1). It is evident that the unhydrolyzed state (state I, SI) exhibits a lower free energy by 2.8 kcal/mol compared to the hydrolyzed state (state II, SII), indicating its higher chemical stability.

The enthalpy of hydrolysis turned out to be 2.8 ± 0.2kcal/mol, which is in perfect agreement with the value resulted from the Umbrella Sampling analysis. Therefore, the entropy change upon reaction is close to zero, and enthalpy is the main driving force in this process.

Considering the chemical reaction alone would not be enough to get an adequate representation of the existing equilibrium. Cys162 discoordination caused the N-loop to become disconnected from the protein’s core, potentially increasing the mobility of this subdomain. Consequently, it could lead to growth in configurational entropy, resulting in a decrease in the relative free energy of SII. On the other side, the configurational effect is usually interfered with by solvation entropy reduction arising from the growth of solvent-accessible area upon partial unfolding [53]. Thus, taking into account solvation and conformation thermodynamics is essential for obtaining precise and comprehensive insights about the relative stabilities of corresponding states.

Classical molecular dynamics simulations (see “Models and Methods”) revealed that the lifetime of the hydrolyzed state in the native configuration does not exceed 20-30 ns (Fig. 4). The system primarily exists in two conformations, with a ratio close to 4:1. In the first one, the N-loop is fully exposed to the solvent and undergoes high-amplitude oscillations. In the other, the loop’s terminus is closer to the protein’s core, and oscillations are predominately absent. Noteworthy, the apo-protein, being not bound to any metal, maintains its original structure even throughout 3000 ns simulations, as indicated by the blue line on the RMSD plot (Fig. 4).

**Figure 4.**
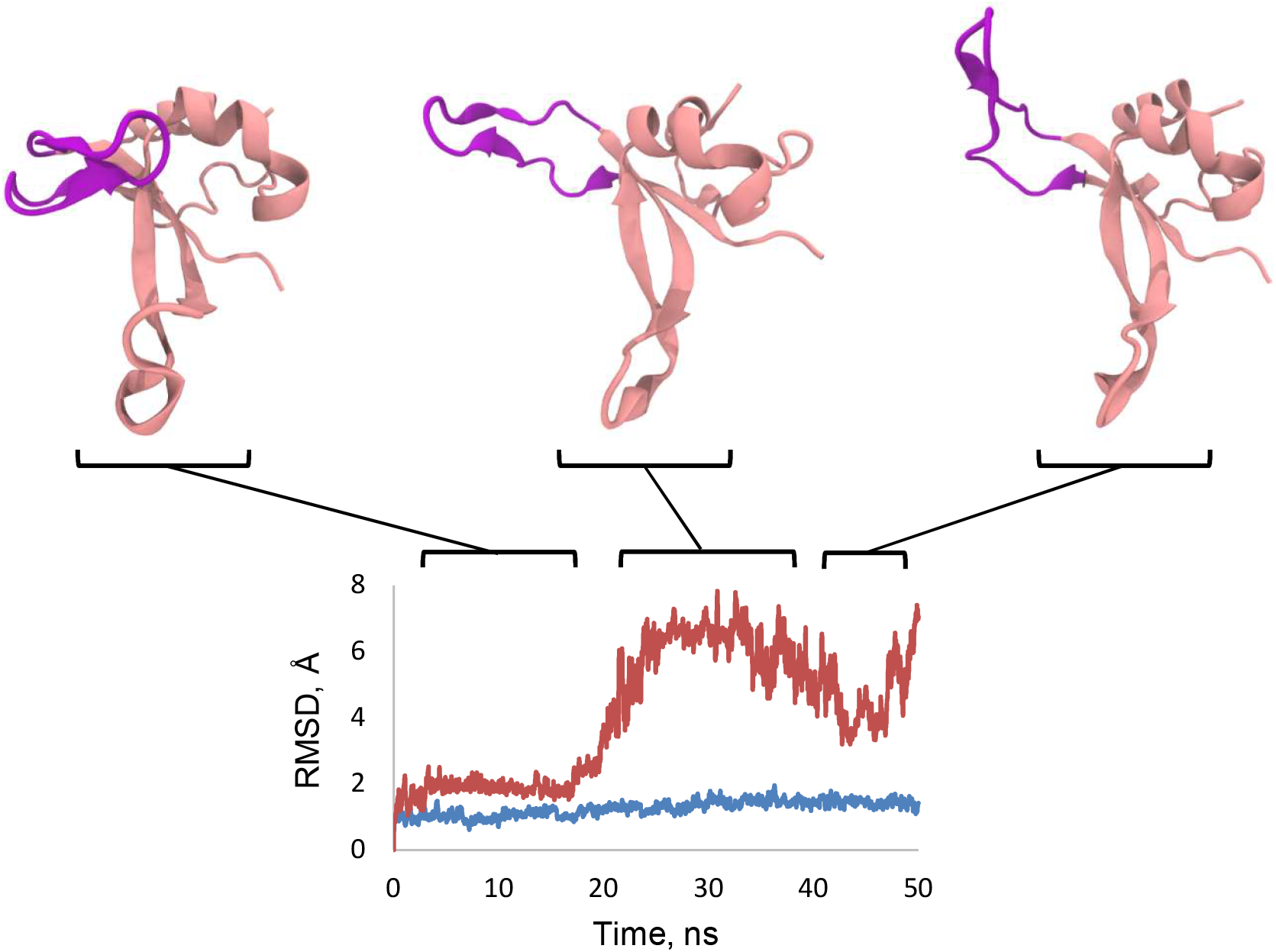
Conformational dynamics of hydrolyzed complex (state II, red color on RMSD plot) and apo-enzyme (blue). N-loop is colored in purple

The total free energy difference between SII and SI has the following expression:

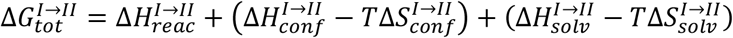

Here, 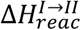 stands for hydrolysis enthalpy, which we showed to be equal to the reaction free energy. In the first brackets, configurational terms are written – differences of intraprotein energy and conformational entropy. The second brackets account for solvation thermodynamics. 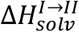 describes how water-water and water-solute interactions energies change upon transition, and 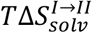 shows the difference of water translational and rotational entropies between SII and SI. A detailed description of corresponding computational protocols is presented in “Models and Methods” section.

Table S2 in supplementary materials contains the final computed values of 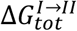 components. One can see the sum of configurational and solvent enthalpies is close to zero, which sends us back to the known fact that protein folding is mainly driven by entropic factor [54]. Excluding hydrolysis energy, the free energy difference between states II and I is equal to -3.4 kcal/mol. This means that in the equilibrium mixture, the fraction of SI will be less than 0.5%. As we mentioned, after being destroyed in a few dozen of ns, the initial state does not regenerate even after 10^3^ ns-long simulations. Given that only the native zinc finger’s fold is supposed to be able to bind nucleic acids, this would necessarily result in domain’s malfunction. Nevertheless, the unfavorability of hydrolysis increases the final free energy difference to -0.6 kcal/mol, which might not be enough to break the functionality of all Zn2 instances. Firstly, upon accounting for computational error even a sign of the free energy difference could change, and, secondly, we will get an equilibrium where comparable fractions of both states coexist, anyway. Therefore, we attempted to search for additional structural and chemical transformations that occur after hydrolysis and govern the stabilization of the protein’s “broken” state.

### Binding site reorganization

We thought that the association of the hydrolyzed uranyl complex with one of the potential metal-binding surfaces found (Fig. 2A) might firm a non-functional state. Out of the two potential binding sites explored, only those composed of Glu190 and Asp191 residues showed an ability to interact with the N-loop. Even without the coordination bond between uranium and acidic cluster modeled, the corresponding loop conformation (Fig. 5) remained intact in 10^3^ ns long molecular dynamics simulations.

**Figure 5.**
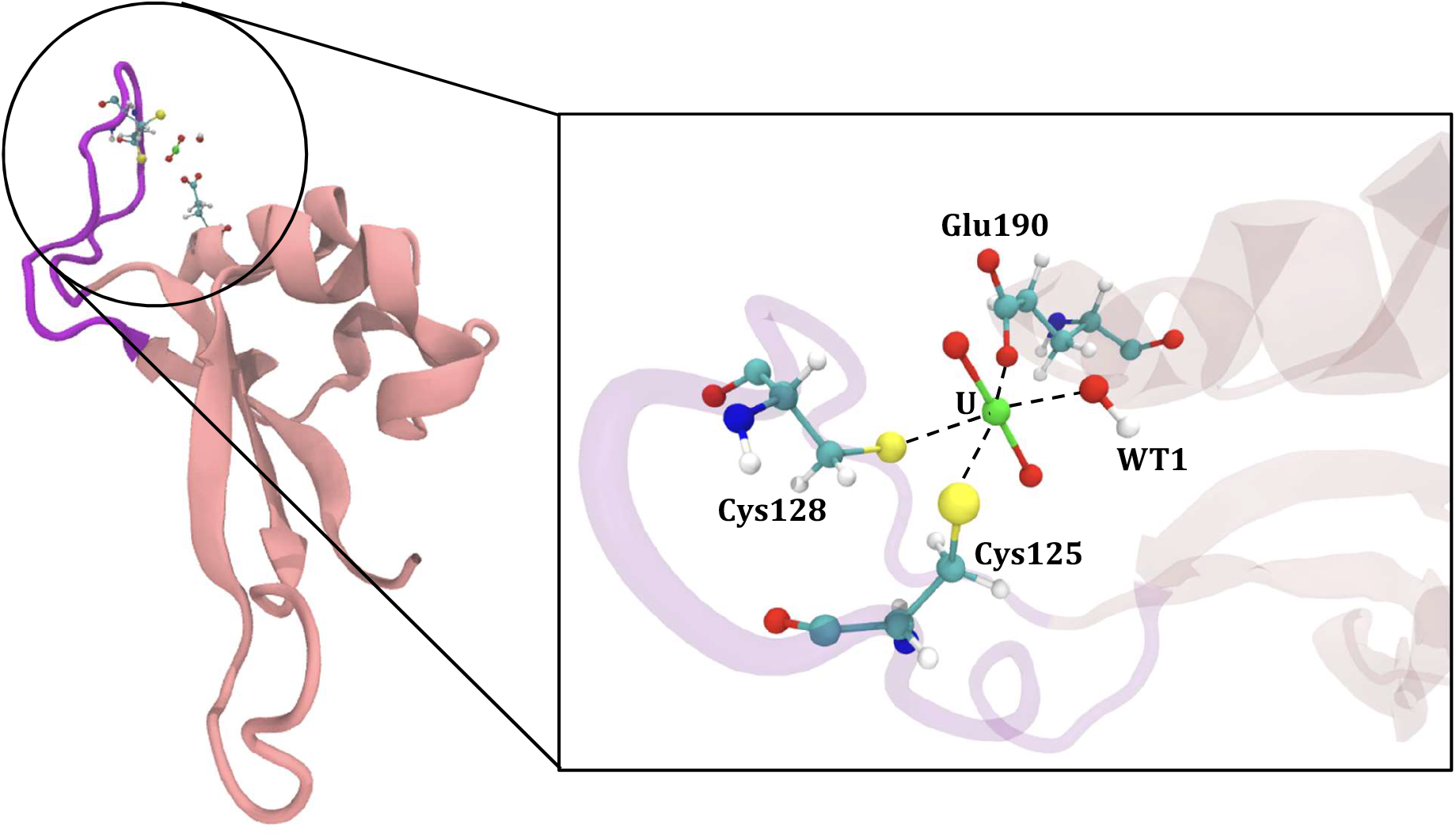
Zinc finger’s conformation and binding site organization in state III (SIII).

Our MD simulations also showed that only Glu190 possessed optimal orientation to associate with uranyl ion. To come up with a possible reaction mechanism, we addressed the stability of uranyl complexes with two methylthiolate groups, one acetate ion and a various number of water molecules and hydroxyl anions. We found out that five-coordinated complexes, as well as those having carboxylate as a bidentate ligand, are energetically and structurally unfavorable. Namely, the U-S bond order in such complexes turned out to be less than 0.1, which corresponds to the absence of coordination bonding. Therefore, it was most likely that the association would proceed by substitution of one of the existing ligands, probably, water:

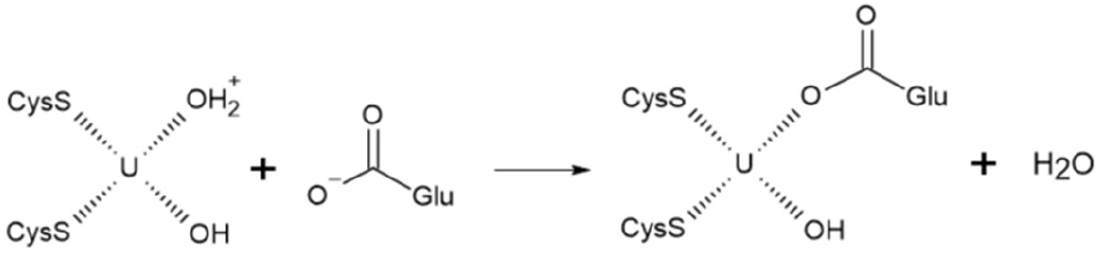

Furthermore, we cannot be certain whether the hydroxyl anion formed in the coordination sphere upon hydrolysis remained deprotonated. However, the association with the carboxylate group of hydroxyl-containing complex [UO_2_(SCH_3_)_2_(OH)(H_2_O)]^−^ would be less favorable than of [UO_2_(SCH_3_)_2_(H_2_O)_2_], so that, the energy of reaction involving the first of them would be a top estimate.

According to our calculations, in the final state (Fig. 5), the bond between uranium and sulfur has a length of 2.77 Å and a bond order of 0.51. In its turn, the coordination bond between uranium and carboxyl oxygen of Glu190 is relatively short (2.31 Å), but the bond order indicates that it is not very strong, with a value of 0.59. However, if we expect that OH-group is going to be reprotonated, the U-SCys bond order will increase to 0.68, and U-OGlu – to 0.88. Therefore, from the chemical point of view, the final complex (SIII) should be more stable than the initial (SI).

As previously, we calculated both free energy change along the association reaction pathway (Fig. S2) and corresponding enthalpy alone. This time, the energy of the product became much lower than that of the reactant. Noteworthy, its absolute magnitude exceeds the energy of internal hydrolysis, making state III chemically more stable than any other. However, in contrast to the internal hydrolysis reaction, the association free energy (−12.1 kcal/mol) turned out to be lower than its enthalpy (−11.7 ± 0.3 kcal/mol). This indicates a non-zero contribution of the entropic term. A water molecule that is released upon reaction, acquires mobility inherent to bulk solvent. Taking into account that QM/MM molecular dynamics simulation covers a short time period and Umbrella Sampling imposes restrictions on translational movements of perturbed molecules, we tended to assume that the 0.4 kcal/mol difference arises from the increase of rotational entropy of water. Bulk TIP3P water has translational 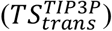 and rotational 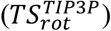 entropies equal to 4.2 and 1.0 kcal/mol, respectively (T = 298 K) [55]. Therefore, the growth of rotational mobility upon association was ∼ 50%.

As for translational entropy, it is non-zero for coordinated water, which is still relatively loosely coupled to uranium (U-H_2_O bond order for SI and SII is 0.26 and 0.25, respectively) and serially being replaced by other solvent waters. However, we are not able to see these replacements in simulations and, consequently, could not estimate this term for our case. Therefore, we can just state that water translational entropy increase 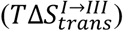 upon association is close to 4.2 kcal/mol.

For SI → SIII transition, the formula for total free energy difference is as follows:

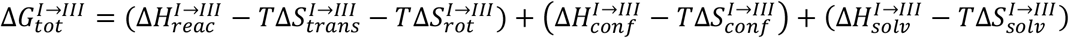

These terms are either similar to those for SI → SII transition or were discussed explicitly. Table S3 contains computed values and shows that 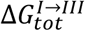 is expected to be close to -7.4 kcal/mol, indicating significant favorability of the “broken” state in comparison to functional one. In state III, the orientation of the N-loop is far away from optimal for effective DNA recognition, so we can be sure that the protein’s DNA-binding interface is now destroyed. Thus, exposure to uranium salts of PARP-like zinc fingers causes irreversible tertiary structure transformations that impair their native functionality.

In our investigation of the chemical and conformational transformations occurring upon exposure of the zinc finger of PARP-1 protein to the uranyl ion, we proceed only from the fact that the loss of DNA-binding activity and zinc depletion are observed under such conditions. However, the mechanism we proposed aligns with additional experimental evidences reported previously. We have demonstrated that the complex formed between the zinc finger and uranyl undergoes significant alterations in its tertiary structure, while the apo-protein, devoid of metal binding, maintains its native conformation over time [11]. Moreover, our mechanistic model reconciles the apparent discrepancy between the findings of Zhou et al., who observed a lack of thermodynamic advantage for uranyl over zinc *in vitro* [11], and Cooper et al., who reported that uranyl replaces zinc from zinc finger *in vivo* [9]. Specifically, Zhou et al. investigated a peptide lacking alternative binding sites, whereas our study revealed that the dissociation of the N-loop alone does not yield significant thermodynamic favorability. However, the subsequent association of uranyl with the missing in peptide Glu190/Asp191 cluster stabilizes the broken state as a whole and reinforces the uranyl-protein interaction.

## Conclusion

By combining different computational chemistry techniques we proposed a molecular model of structural changes occurring in PARP-like zinc fingers upon their exposure to uranium salts. We showed that the binding of zinc and uranyl ions is competitive: the latter occupies the metal-affine site that is partially composed of amino acid residues (Cys125, Cys128) that bind zinc in natural (functional) conformational state. This is consistent with experimental studies reporting that the interaction of zinc finger with uranyl ion results in zinc loss. However, the uranyl ion complex in the native metal-binding site turned out to be unstable. U-Cys162 coordination bond undergoes spontaneous hydrolysis leading to the destruction of the DNA-binding interface, which was also observed in *in vitro* experiments. Further structural reorganizations condition uranyl ion association with the carboxyl group of Glu190, so that the final uranyl-binding site is formed by Cys125, Cys128, Glu190 residues and one water molecule (or hydroxyl anion). The free energy difference between the “broken” and functional states of the protein-uranyl complex is close to -7.4 kcal/mol, which corresponds to near to irreversible transformation.

## Acknowledgments

The authors acknowledge the use of the Skoltech HPC cluster Zhores for obtaining the results presented in this paper [56]. The authors are also grateful to Marina Pak and Natalia Sivitskaya for assisting while preparing the manuscript.

