## Supplementary Material for "*In silico* simulations reveal molecular mechanism of uranyl ion toxicity towards DNA-binding domain of PARP-1 protein"

### SUPPLEMENTARY MATERIALS

**Table S1.** Bonding properties of uranyl complexes with different ligand compositions. N stands for a number of ligands of a particular type. For those formulas that have several structural representations, average values are presented.

| *Formula* | *N($CH_3S^-$)* | *N($H_2O$)* | *N($OH^-$)* | ***Bond lengths*** | | | ***Mayer bond orders*** | | | ***IBSI*** | | | ***Charge trans. to $UO_2$ (-e)*** | | |
|---|---|---|---|---|---|---|---|---|---|---|---|---|---|---|---|
| | | | | *U-$SCH_3$* | *U-$H_2O$* | *U-OH* | *U-$SCH_3$* | *U-$H_2O$* | *U-OH* | *U-$SCH_3$* | *U-$H_2O$* | *U-OH* | *$SCH_3$* | *$H_2O$* | *OH* |
| ***s2w2*** | 2 | 2 | 0 | 2.67 | 2.48 | x | 0.85 | 0.27 | x | 0.26 | 0.21 | x | 0.63 | 0.16 | x |
| ***s2w1oh1*** | 2 | 1 | 1 | 2.73 | 2.56 | 2.18 | 0.79 | 0.25 | 0.56 | 0.23 | 0.18 | 0.42 | 0.55 | 0.14 | 0.37 |
| ***s2oh2*** | 2 | 0 | 2 | 2.82 | x | 2.22 | 0.65 | x | 0.44 | 0.2 | x | 0.38 | 0.47 | x | 0.34 |
| ***s3w1*** | 3 | 1 | 0 | 2.70 | 2.53 | x | 0.81 | 0.24 | x | 0.24 | 0.19 | x | 0.56 | 0.16 | x |
| ***s3oh1*** | 3 | 0 | 1 | 2.78 | x | 2.2 | 0.49 | x | 0.5 | 0.21 | x | 0.4 | 0.38 | x | 0.34 |
| ***s2w3*** | 2 | 3 | 0 | 2.71 | 2.59 | x | 0.87 | 0.25 | x | 0.24 | 0.17 | x | 0.62 | 0.19 | x |
| ***s2w2oh1*** | 2 | 2 | 1 | 2.75 | 2.64 | 2.22 | 0.66 | 0.25 | 0.60 | 0.19 | 0.15 | 0.37 | 0.5 | 0.18 | 0.4 |
| ***s2oh3*** | 2 | 0 | 3 | 3.06 | x | 2.29 | 0.44 | x | 0.51 | 0.13 | x | 0.35 | 0.25 | x | 0.34 |
| ***s3w2*** | 3 | 2 | 0 | 2.70 | 2.7 | x | 0.78 | 0.26 | x | 0.2 | 0.15 | x | 0.69 | 0.13 | x |
| ***s3oh2*** | 3 | 0 | 2 | 3.02 | x | 2.28 | 0.36 | x | 0.47 | 0.14 | x | 0.33 | 0.26 | x | 0.41 |

**Table S2.** Solute and solvent thermodynamic properties change upon transition from state I to state II.

| *Free energy component* | *Value, kcal/mol* |
|---|---|
| $\Delta H_{reac}^{I\to II}$ | 2.8 |
| $\Delta H_{conf}^{I\to II}$ | 4.4 |
| $T\Delta S_{conf}^{I\to II}$ | 19.4 |
| $\Delta H_{solv}^{I\to II}$ | -4.1 |
| $T\Delta S_{solv}^{I\to II}$ | -15.7 |
| $\boldsymbol{\Delta G_{tot}^{I\to II} = -0.6\ kcal/mol}$ | |

**Table S3.** Solute and solvent thermodynamic properties change upon transition from state I to state III.

| *Free energy component* | *Value, kcal/mol* |
|---|---|
| $\Delta H_{reac}^{I\to III}$ | -8.9 |
| $\Delta H_{conf}^{I\to III}$ | 1.8 |
| $T\Delta S_{conf}^{I\to III}$ | 1.1 |
| $T\Delta S_{rot}^{I\to III}$ | 0.4 |
| $T\Delta S_{trans}^{I\to III}$ | < 4.2 |
| $\Delta H_{solv}^{I\to III}$ | -2.1 |
| $T\Delta S_{solv}^{I\to III}$ | -7.5 |
| $\boldsymbol{\Delta G_{tot}^{I\to III} \sim -7.4\ kcal/mol}$ | |

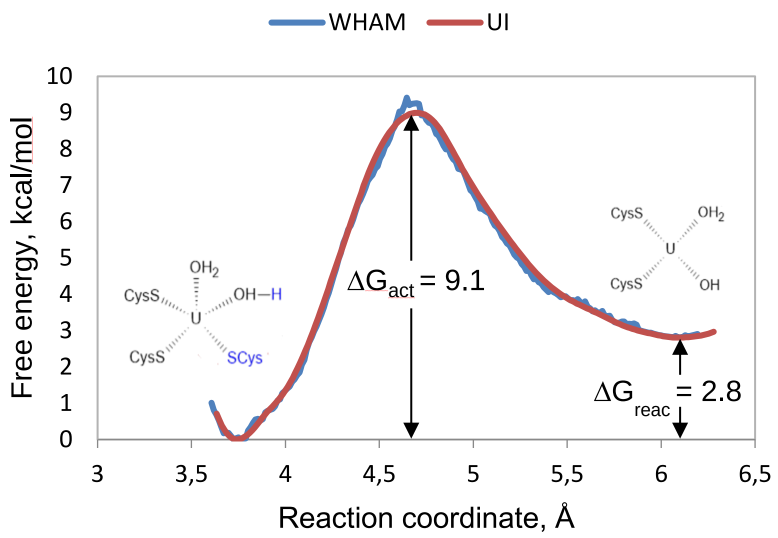


**Figure S1.** Internal hydrolysis reaction free energy profile derived from Umbrella sampling calculations. Color corresponds to US results analysis method. The reaction coordinate is d(OH2$^{WT1}$, H2$^{WT1}$) + d(U$^{UO2}$, SG$^{Cys162}$).

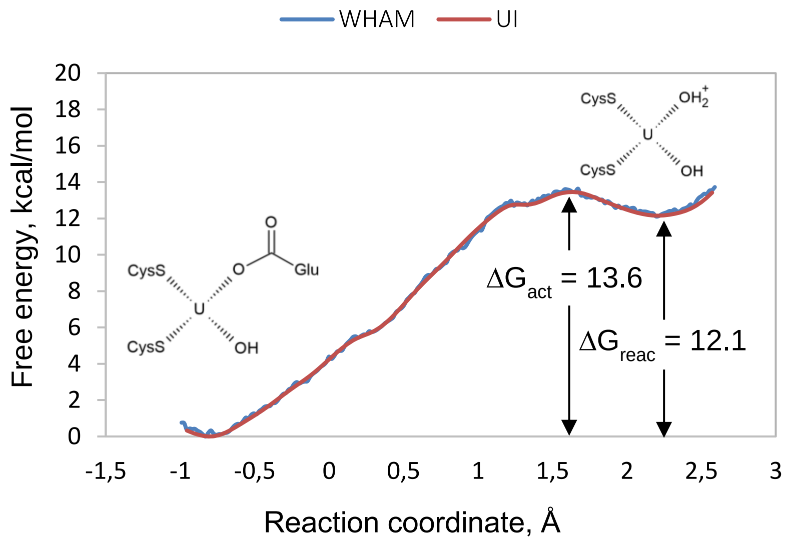


**Figure S2.** Association reaction free energy profile derived from Umbrella sampling calculations. Color corresponds to US results analysis method. The reaction coordinate is d(U$^{UO2}$, COO$^{Glu190}$) – d(OH2$^{WT2}$, U$^{UO2}$).
